# A scalable assay for chemical preference of small freshwater fish

**DOI:** 10.1101/2022.07.10.499464

**Authors:** Benjamin Gallois, Léa-lætitia Pontani, Georges Debrégeas, Raphaël Candelier

**Author notes:** Correspondence: Raphaël Candelier.

## Abstract

Sensing the chemical world is of primary importance for aquatic organisms, and small freshwater fish are increasingly used in toxicology, ethology, and neuroscience by virtue of their ease of manipulation, tissue imaging amenability, and genetic tractability. However, precise behavioral analyses are generally challenging to perform due to the lack of knowledge of what chemical the fish are exposed to at any given moment. Here we developed a behavioral assay and a specific infrared dye to probe the preference of young zebrafish for virtually any compound. We found that the innate aversion of zebrafish to citric acid is not mediated by modulation of the swim but rather by immediate avoidance reactions when the product is sensed and that the preference of juvenile zebrafish for ATP changes from repulsion to attraction during successive exposures. We propose an information-based behavioral model for which an exploration index emerges as a relevant behavioral descriptor, complementary to the standard preference index. Our setup features a high versatility in protocols and is automatic and scalable, which paves the way for high-throughput preference compound screening at different ages.

## 1 INTRODUCTION

Assessing the attraction or repulsion of fish for chemical compounds has both direct economic implications - as chemical lures are commonly used in industrial and recreational fishing (Hara, 1992) - and a strong practical importance for several fields of research. In toxicology, fish are used to study the effects of alcohol and drug abuse both at the behavioral and neuronal scales (Stewart et al., 2011). In ecotoxicology, environmental contaminants can be heterogeneously distributed in space (Steele et al., 1990; Brady et al., 2017) and their impact on an ecosystem can greatly differ depending on whether animals tend to seek or avoid regions of high concentration (Höglund, 1961; Jacquin et al., 2020). In neuroscience, attractive compounds are often used as rewards in associative learning paradigms (Pearce and Bouton, 2001), though robust attractants for aquatic model systems like zebrafish (*Danio rerio*) and, more recently *Danionella Cerebrum*, are still largely unknown. In addition, the variable responses to chemicals across fish species (Hara, 1992; Kasumyan and Doving, 2003), even evolutionary close, and the plethora of possible compounds to test call for standardized, high-throughput behavioral assays for chemical preference adapted to fish.

During the last two decades, the use of small aquatic animals for toxicity screenings has steadily increased. Zebrafish is at the forefront of this effort (MacRae and Peterson, 2015; Kim et al., 2020) due to its small size, high fecundity, permeability to small molecules, transparency at the larval stage, and early development of dopaminergic projections in the forebrain forming a highly conserved pathway analog to the mesolimbic system of mammals (Stewart et al., 2011). Automated platforms manipulating live animals back and forth from multi-well plates to microscopes have led to high-throughput imaging of immobilized larvae (Pardo-Martin et al., 2010, 2013). Still, assessing toxicity only requires two *endpoints* measures before and after exposure (Horzmann and Freeman, 2018), while assessing preference is way more challenging: it demands monitoring freely moving animals having a choice to favor or avoid a compound.

Quantifying the chemical preference of fish can be done either by direct *place preference* (PP) or *conditioned place preference* (CPP) assay. The latter is a type of Pavlovian conditioning during which the fish first associates a chemical stimulus with a conditioned stimulus of another modality (*e*.*g*. color, visual pattern) and then reveals its preference to regions where the conditioned stimulus is localized, in the absence of chemical cues (Tzschentke, 2007; Mathur et al., 2011; Collier and Echevarria, 2013; Blaser and Vira, 2014). It is particularly well-suited for adult fish since it eliminates all hydromechanical issues associated with chemical transport in the test chamber, but it cannot be employed with young fish since it relies on the association and memorization of different cues. Besides, the CPP experiments are days to weeks long and entail numerous fish manipulations, so they are bound to low-throughput assays.

For larvae and juvenile fish, the place preference paradigm remains the only option and turns out to be experimentally challenging for in-depth behavioral analyses or medium to high-throughput screening. Several geometries have been used for different purposes: the setup of (Herrera et al., 2021) relies exclusively on diffusion to create a gradient of concentration but has been limited so far to salt ions that have a high diffusion coefficient (Nielsen et al., 1952). With organic molecules, it would take from 2 to 10 times longer to establish a stable gradient. In Bretaud et al. (2007) the fish evolve freely in two facing counterflows with a midline drain, which ensures a sharp and self-healing demarcation between the product and buffer sides; however, the behavioral reaction to chemical information at the interface may be influenced by the concurrent change in flow direction. The U-shaped *choice flume* geometry proposed in Mann et al. (2003) and Hussain et al. (2013) resolves this issue by generating adjacent coflows in the same direction, but the physical separation present in a large portion of the test chamber limits the moments where the fish can actually obtain information over the chemical field; this leads to unnecessarily long and fish-exhausting runs.

Here, we opted for the geometry previously employed for much larger fish in the *fluvarium* of Kroon and Housefield (2003), with a strong emphasis on automation and scalability and drastic simplifications in flow preparation allowed by the millimetric size of the chamber. We developed a bio-compatible infrared dye to simultaneously monitor the trajectories of small fish and the concentration field. The setup is open-source, low-cost, and versatile as it offers a large variety of possible protocols with limited fish manipulation.

We first review the universal hydromechanical limitations inherent to preference assays at all zebrafish developmental stages, and then we present preference results for young zebrafish the in presence of citric acid and ATP. We found that citric acid is always repulsive above an age-dependent threshold and that in the presence of ATP at high concentration juvenile zebrafish switch from repulsion to attraction after a few exposures. We propose an information-based model that allows us to define preference and exploration as independent behavioral measurements and provides a thorough description of all observed behavior.

## 2 MATERIALS AND METHODS

### 2.1 Animals

Experiments were performed on wild-type zebrafish (*Danio rerio*) aged 6 to 31 days post-fertilization (dpf). Larvae were reared in Petri dishes in E3 solution on a 14/10 hours light/dark cycle at 28°C and were fed powdered nursery food every day from 6 dpf. After 11 days, fish were transferred to tanks and fed powdered nursery food and rotifers. *Danionella Cerebrum* (DT) adults were grown at a water temperature of 25 to 26°C, pH of 7.1 to 8,1, and a conductivity of 300 to 450μS. Adult DT are fed with a mix of Gemma Micro 75 and 150 (Skretting, USA) ang algae, twice a day.

All experiments were approved by *Le Comité d’Éthique pour l’Expérimentation Animale Charles Darwin* C2EA-05 (02601.01).

### 2.2 Millifluidic chip and microvalve manifolds

The millifluidic chip was made by assembling two laser-cutted (Full Spectrum Laser, PS20), 3mm or 5mm-thick PMMA slabs, bonded together under press with 70% acetic acid at 65°C for 1 hour. The chip total dimensions were 130×80mm and the chamber size was 60×30mm. Custom 3D-printed connectors in PETG (Prusa MK3) held the inlet and outlet tubes. 40mm-long tuyeres placed before and after the chamber were designed to create a laminar flow. Two rigid grids, first 3D-printed in PETG and then drilled with 280 holes of diameter 500μm (Minitech Mini-Mill/GX) maintained the fish in the chamber. In addition, the chamber was covered by a 2mm-thick PMMA laser-cutted slab.

The flow rate in the chamber was fixed at 60μL s^−1^. Six 3-way microvalves (The Lee Company, LHDA0531115H) were used to control which solution syringes were filled with, and to which side of the chamber they infused (see manifold logic in supplementary figure 1). Two custom manifolds supporting the microvalves and ensuring a strong fitting of the tubes were 3D-printed in PETG.

### 2.3 Dye

We used a diluted emulsion of silicone oil droplets as a dye during preference protocols. For emulsion preparation, 10mL of silicon oil (silicone 350cst, Sigma Aldrich) is first coarsely emulsified in 22mL of a viscous aqueous buffer containing 10mM Sodium Dodecyl Sulfate (SDS) and 1 wt% alginate (Alginic acid sodium salt from brown algae, Sigma Aldrich). This crude emulsion is then injected into a narrow gap couette mixer, with a gap size of 100μm, and sheared at 30rpm. The resulting emulsion is washed twice in 250mL of aqueous solutions of 8 to 10mM SDS in a separating funnel. The top phase containing the dense cream is finally collected. The droplets display an average diameter of 5μm with a polydispersity of ∼15%. Alternatively, similar droplets can be obtained through pressure emulsification. In that case, 5mL of silicone oil (silicone 50cst, from Sigma Aldrich) is extruded through a Shirasu Porous Glass (SPG) membrane decorated with pores of 1μm diameter, using a membrane emulsification device (Internal pressure type micro kit, SPG Technology Co., Ltd). The droplets are directly extruded into a vigorously agitated aqueous solution containing 10mM SDS. In this case the droplets also yield an average diameter of 5μm, with a polydispersity of ∼10%. All emulsions are stable during several months at room temperature.

### 2.4 Illumination

The setup has two possible illumination pathways and can work either in bright field or dark field mode (supplementary figure 2). In bright field, a diffuser (translucent paper) is placed at mid-height of the illumination box and the walls are non-reflective. In dark field, direct transmission from the LED is suppressed by an opaque blocker while reflections on the walls are permitted, such that light arrives obliquely on the millifluidic chamber.

Examples with both illuminations are given in supplementary movie 1. The appropriate illumination mode depends on the species, age and pigmentation of the animal. As a general rule of thumb, bright field is better suited to opaque animals (*e*.*g*. juvenile zebrafish) while the dark field is better suited to transparent animals (*e*.*g*. DT larvae).

### 2.5 Image processing and trajectory analysis

For all movies, the position of the animals’ head was extracted automatically using FastTrack (Gallois and Candelier, 2021). The concentration maps were computed with Matlab custom scripts.

Swim bouts were determined with the same method as in Olive et al. (2016). The time-based preference index was calculated using the time spent in each zone, defined by the mean interface position. For the events-based analysis, engagements and withdrawals at the interface were visually counted in a blinded protocol such that the person counting the events does not know either what is the product or its concentration, or if there is no product at all in the dyed region (control). Each movie has been reviewed by two independent persons, and a check for inconsistencies pointed out a few mistakes, that were corrected. The counted events have then been pooled together to get an average value. When specified, the given number of fish (*n*) stands for the number of trajectories per analysis.

## 3 RESULTS

### 3.1 Setup

#### 3.1.1 General design

The setup is an open-source assay designed for the high-throughput evaluation of chemical preference. We packed all mechanical (illumination box, syringe pump, valve manifold) and electrical (power supply, computer, valve controller) parts in a vertical structure (figure 1A) with a low table-top footprint (25×35cm). All components were selected to serve their function robustly and accurately for a low price, such that the complete bill of materials remains below 2k€ (supplementary table 1). It is thus particularly easy to build several instances that can be run in parallel in a reduced space and facilitate the screening of several chemical compounds, at many concentrations and with animals at different developmental stages.

**Figure 1.**
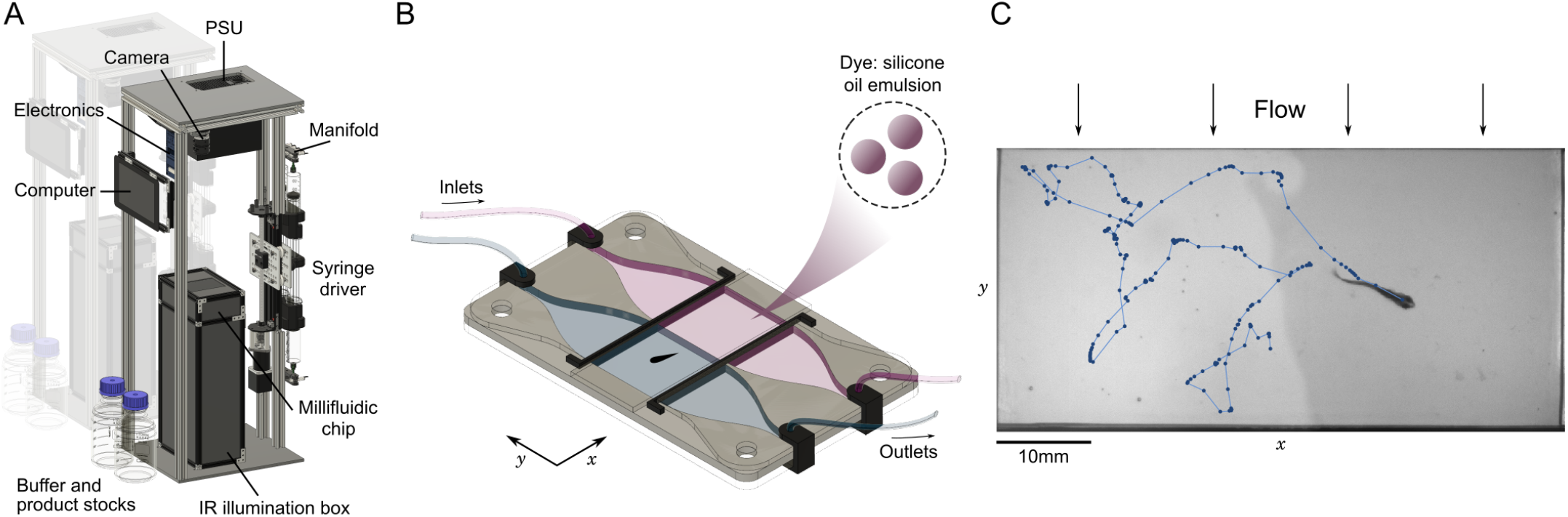
Automated assay for preference in fish. **(A)** Annotated view of the rig. The vertical structure has a low table-top footprint, allowing to align multiple setups side-by-side. **(B)** Schematic view of the millifluidic chamber during dual infusion of buffer (water) on one side, and a mixture of a water-soluble compound and a diluted emulsion of silicone oil droplets used as a dye on the other side, creating an interface at the center of the imaging chamber. **(C)** Sample image and trajectory over 25s of a juvenile zebrafish (28 dpf) swimming in the chamber. Due to the mild transverse flow, the interface continuously regenerates in less than 10 seconds.

At the core of the setup, a millifluidic chip hosts the imaging chamber where animals freely evolve in two co-flowing water-based fluids, with an optional interface at the center consisting of a steep chemical gradient. All freshwater animals below 5mm in height can fit in the chamber, including zebrafish larvae, fry and juveniles (Singleman and Holtzman, 2014) (figure 1C) and DT at any age (Schulze et al., 2018; Rajan et al., 2021) (supplementary movie 1). Multiple fish can be imaged simultaneously in the chamber as long as they are small enough and do not interact with each other, as is the case at the larval stage. We have checked that in the absence of product and dye both larval and juvenile zebrafish do not have a preferred side of the chamber (supplementary figure 3). We also tried to record the behavior of *Xenopus tropicalis* tadpoles, but they remained much more passive than fish and rarely crossed the interface (data not processed).

#### 3.1.2 Automation and versatility

We identified four canonical protocols that can be implemented in the setup (supplementary figure 4) with human intervention being essentially limited to preparing the solutions and inserting/removing the animal from the chamber. First, successive exposures can be applied with different durations. For long exposure times, however, fast refills of the syringes are necessary every 15min of infusion. Second, a stock solution can be diluted at will during the syringes filling phase (by filling up to a given volume and completing with buffer) enabling successive exposures with controlled concentrations. Third, different compounds can be tested sequentially. In this case, a human operator has to switch the tube inlets to the correct stock solution before each syringe refill, though an automated switching module could be easily developed based on the same manifold system. Finally, two solutions can be infused simultaneously on both sides of the chamber, typically a compound to test and a dye on one side and buffer on the other side to create a preference assay.

Though we restricted ourselves to a simple preference-testing protocol with two exposures per fish in this study, these four canonical protocols can be mixed at will to create a very large panel of automated behavioral assays.

#### 3.1.3 Infrared dye

In many previous experimental studies exploiting direct chemical place preference (Braubach et al., 2009; Koide et al., 2009; Hussain et al., 2013; Krishnan et al., 2014; Wakisaka et al., 2017; Lucon-Xiccato et al., 2020; Choi et al., 2021) the tested compound was not marked, inserted locally with a brief shot and was assumed to remain localized while the animal could freely explore the test chamber. Hence there have been large uncertainties over when the animal was actually in contact with the compound and at what concentration, which restricted the behavioral analyses to basic measurements over short runs. Moreover, as we shall see in section 3.2, the initial concentration fields are rapidly altered and without renewal these uncertainties grow at a pace comparable to the fish dynamics.

The concentration field of the tested compound could be easily obtained by mixing it with a dye and imaging with the same camera monitoring the trajectory, but finding an appropriate dye revealed to be challenging. First the animal should not see it, otherwise the trajectories could be biased by visual cues like the innate dark avoidance of larval zebrafish, termed *scotophobia* (Steenbergen et al., 2011; Karpenko et al., 2020). We thus used a dark box with near-infrared (IR) illumination that is invisible to the fish. Then, the IR dye should be absorbent enough on a few millimeters, available in large quantities, chemically inert, non-toxic and neutral to the preference of the animal. As we couldn’t find a chemical compound meeting all these requirements, we developed a custom dye consisting of an emulsion of colloidal droplets of silicon oil with a controlled size of 5μm that can be easily replicated (see Materials and Methods). In contrast to chemical compounds that absorb light at a given wavelength, such a “physical” dye scatters light at all wavelengths, including the near-IR domain. Thus the illumination source, filters and camera sensor can be chosen independently of the dye itself.

Given the dilution of the colloidal dye into the experimental medium, the change of viscosity is undetectable by the animal and there are only trace amounts of surfactant (about 1μM). At such concentrations, SDS was shown to have no effect on zebrafish mortality (Ali et al., 2011) and we did not observe any deleterious effect on the fish after the experiments. Moreover, other biocompatible surfactants and oils can be used to prepare the colloidal emulsion, such that it is non-toxic to more delicate aquatic species. We have checked that the dye is neutral to zebrafish in terms of preference index (supplementary figure 5).

### 3.2 The many roles of the flow

Fish gather information over their chemical environment whenever they detect variations in concentration. To accurately monitor when these moments occur and to maximize their effect, it is experimentally desirable to keep a sharp contrast between regions with and without the tested product and, conversely, to avoid zones of mixing. In the general case, mixing is caused by a combination of diffusion at short length scales and advection over longer distances. We quantify these effects in this section, and show that setting a mild flow has three distinct benefits: maintaining a sharp interface, homogenize the dynamics of larvae and healing motion-induced mixing for larger fish.

#### 3.2.1 Diffusion at the interface and larvae’s motion

Molecular diffusion is the unavoidable and irreversible process by which thermal motion spreads the molecules in all available space. The timing of this process is controlled by a diffusion coefficient *D* appearing in the first Fick’s law, which depends on the molecule, solvent and temperature. The tables of Yaws (2007) list the diffusion coefficient of 5000 organic compounds in water at 25°C. Despite the variety of molecular weights and structures in this database, the values of *D* are tightly packed with a mean value of 6.8 × 10^−10^m^2^ s^−1^ and a standard deviation of 2.2 × 10^−10^m^2^ s^−1^ (see distribution in figure 2), and the extremal values are contained within a single decade. One can thus reasonably expect that any organic molecule used for testing fish chemical sensing has a diffusion coefficient in this range. For instance, citric acid and ATP – the two compounds used later in his study – have *D*_*CA*_ = 6.9 × 10^−10^m^2^ s^−1^ and *D*_*ATP*_ = 7.2 × 10^−10^m^2^ s^−1^ (Bowen and Martin, 1964). Our dye is composed of much larger objects which, according to the Stokes-Einstein equation, yields a diffusion coefficient orders of magnitude below (*D*_*dye*_ ∼ 10^−13^m^2^ s^−1^).

**Figure 2.**
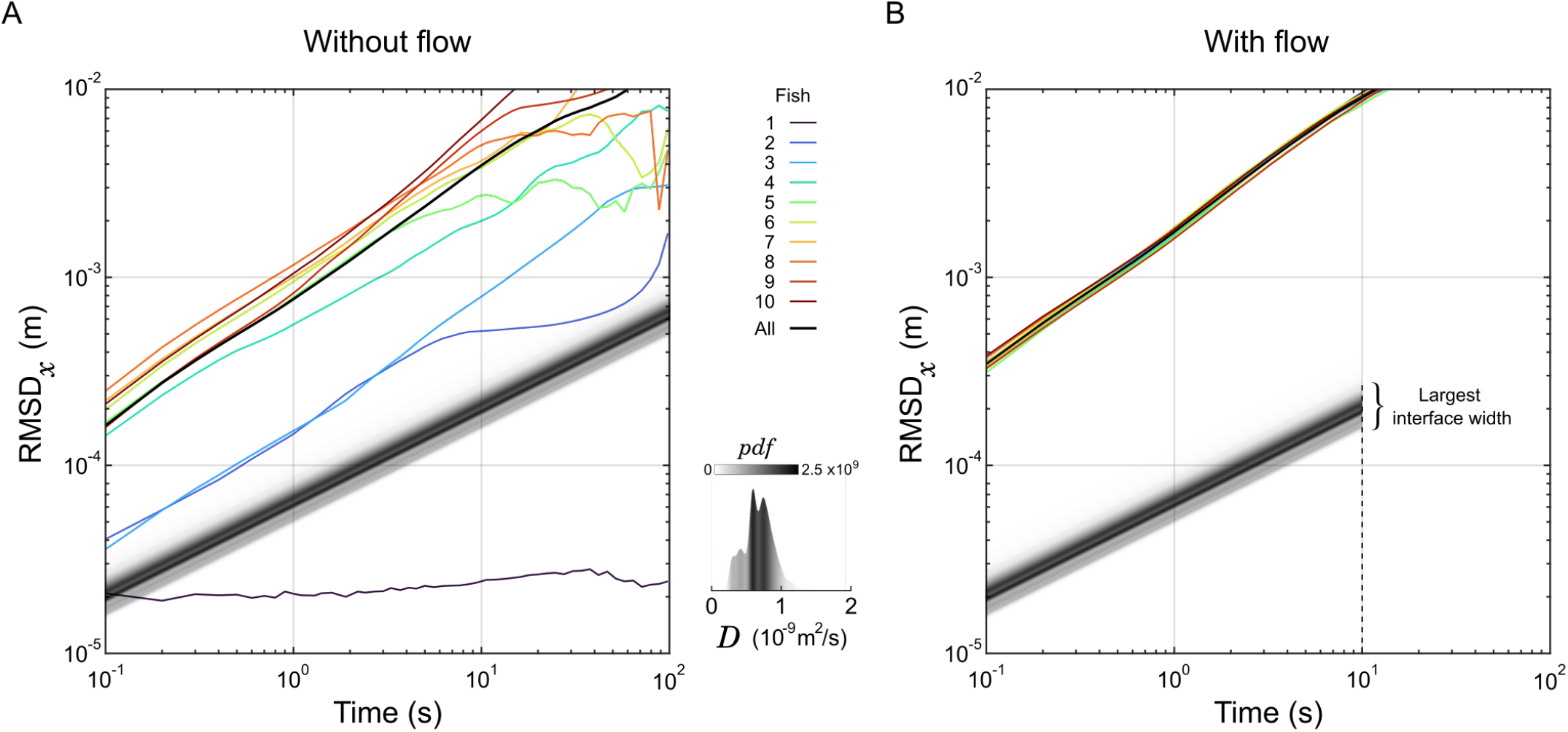
Dynamics of larval zebrafish and diffusion of organic compounds. **(A-B)** The root mean square displacement along the *x*-axis (RMSD_*x*_) of 10 individual larvae are displayed as a function of the sampling duration in the absence (A) or presence (B) of a flow. The RMSD computed over all larvae are displayed in black curves as well. For comparison, the displacement of the front of an interface at a given normalized concentration *α* is a straight line with slope 1/2. These lines are represented for *α* = 1% in shades of grey corresponding to the distribution of the diffusion coefficient *D* of 5000 organic compounds. The latter distribution is also represented separately in the insert. Under flow (B), the largest interface width sits at the end of the test chamber, corresponding to the spreading of a front in 10s.

Given the geometry of our experimental chamber and the constant flow, the concentration profile in the direction perpendicular to the flow (*x*) changes along the flow direction (*y*) exactly like a perfect step of concentration evolves with time in a classical, one-dimensional linear channel geometry. So, assuming an initial condition where the product is confined on one side at concentration *c*_0_ with a perfect step located at the interface in *x*_0_, one can solve the second Fick’s law (El-badrawi et al., 1980) and predict the displacement of the front *d*(*t*) = |*x*(*t*) − *x*_0_|at a given level of normalized concentration *α* = *c/c*_0_, based on the formula:

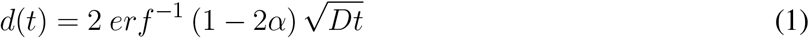

Thus, in logarithmic scale the displacement of a front as a function of time is a straight line with a slope 1/2. We displayed this evolution for all the organic molecules of Yaws’ database at *α* = 1% in figure 2A, with shades of gray corresponding to the distribution of *D*. As a particle of fluid passes through the chamber in 10s, the molecular diffusion is constrained below ∼0.3mm on both sides of the interface mid-position (figure 2B), securing a steep gradient; this is the first role of the flow in our setup.

Then, we tracked the individual trajectories of 10 zebrafish larvae (6 dpf) in the absence of product, dye, and flow, and their root mean squared displacement (RMSD) in one direction as a function of the sampling time are overlaid in figure 2A. Apart from one larva that didn’t move at all, they were faster than the predicted motion of all organic compounds, and on average won the race by approximately one decade. However, there are huge differences in average speed spanning orders of magnitude from fish to fish, that could be due to long resting times largely distributed in frequency and duration.

We then tested the dynamics of the same 10 larvae by setting a flow of buffer in the experimental chamber. The fish were forced to swim against the flow to compensate for the drift (*rheotaxis*) and could not rest. Remarkably, all curves of RMSD in the direction transversal to the flow collapsed on the same line with very little variability (figure 2B). The exploratory dynamics were also largely increased since all fish were traveling faster than the fastest fish in the absence of flow. So, the second role of the flow is to increase and homogenize the exploratory dynamics of the larvae.

#### 3.2.2 Motion-induced mixing with large fish

Larger fish are not concerned by this discussion since they are less prone to freezing and, as they move orders of magnitude faster than larvae, the same flow does not affect their global orientation, hence they do not display rheotactic behavior. However, as the fish size and speed grow, the Reynolds number increases above the transition to turbulence – up to *Re* ∼ 5000 during adult swim (Müller et al., 2000) – and vortices are shed away after each tail beat (Mwaffo et al., 2017; Liao, 2007), creating plumes that can extend far away from the interface (figure 1C, figure 3A-B, supplementary movie 1) and leading to complex eddies in a few successive beats.

**Figure 3.**
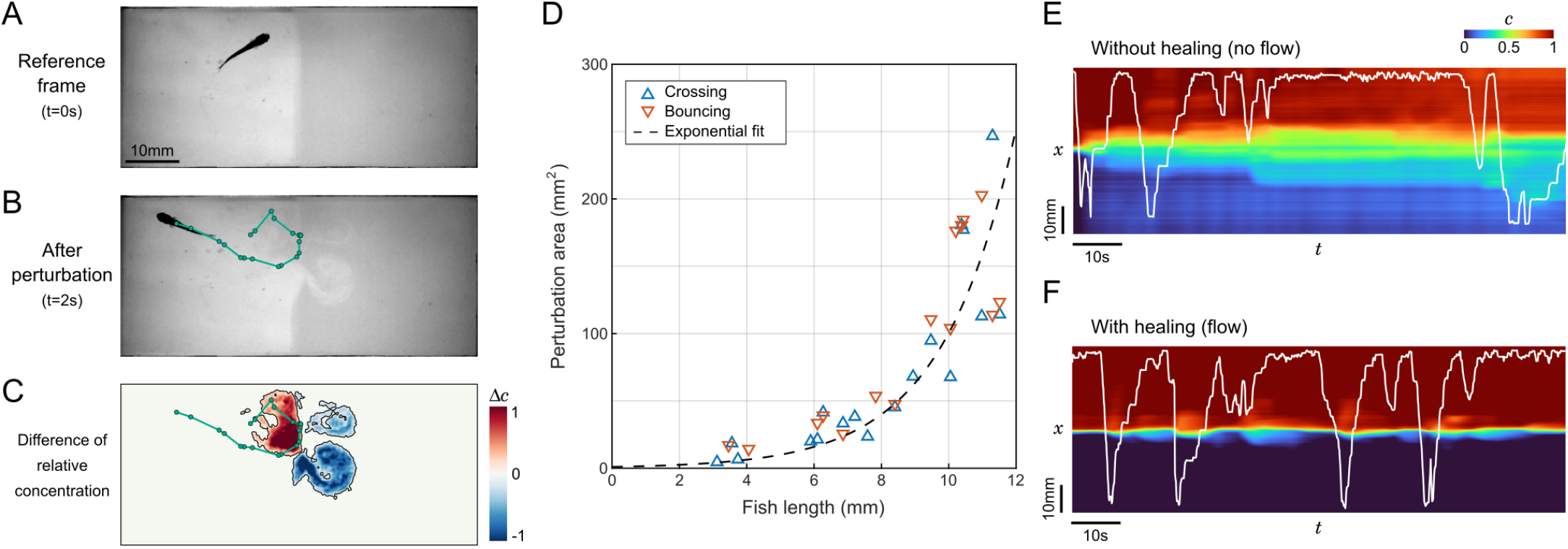
Perturbation of the interface by fish movements. **(A-B)** Sample frames before and after a 2s trajectory of a juvenile zebrafish bouncing on the interface. The trajectory is highlighted in green. **(C)** Field of the difference in relative contentration for the same short trajectory. The perturbed area is contoured in black. **(D)** Perturbation area as a function of fish length for 33 randomly selected events where different fish either passes through the interface (*n*=18) or bounces onto it (*n*=15). **(E-F)** Relative concentration averaged over *y* as a function of time. The position of the same juvenile zebrafish is displayed in white. Without flow, the successive swim movements rapidly spread the compound in the whole chamber (E), while under flow the interface self-heals to keep a controlled separation (F).

Close to the interface, this is a very efficient vector of mixing. Our video recordings can be analyzed to determine the perturbation of the relative concentration field during brief contacts with the interface (figure 3C). These contacts can be separated into two categories: “crosses” when the animal switches sides, and “bounces” when the animal detects the chemical gradient at the interface and ends up staying on the same side. We set a threshold on the difference in normalized dye concentration at 0.2 (figure 3C) and extracted the perturbed area for events generated by fish at various ages (figure 3D). The perturbation area increased rapidly with fish length, without a difference between bounces and crosses. The points were better fitted by an exponential law than with a parabola, indicating that the growth is faster than linear.

To better apprehend how dramatic this mixing can be in a preference assay, we analyzed the degradation of the concentration field in sequences starting with a neat interface where a juvenile fish could freely evolve, with and without flow (figure 3E-F). In the absence of flow, the degradation was very fast since the whole chamber featured a change of concentration of at least 10% in 12s. On longer timescales, the mixing induced by the fish exploration did not lead to a complete homogenization though, and an irregular gradient reminiscent of the initial configuration remained visible. In the presence of flow, the interface self-healed after each perturbation by the fish. In control experiments, the average time spent on one side of the setup, defined as the mean time between two contacts with the interface, was found to be 52s for zebrafish larvae and 9.6s for juveniles, *i*.*e*. larger than the interface self-healing timescale. These timescales confirm that the employed flow rate is high enough to regenerate a sharp concentration profile between successive contacts, at any age.

### 3.3 The repellent effect of citric acid

Let us now turn to two case studies of zebrafish chemical preference, starting with repulsion to a low pH environment. We chose citric acid (CA), a weak acid with low toxicity that has been previously shown to trigger aversive behavioral responses (Candelier et al., 2015).

Representative trajectories of a fish in contact with CA are shown in figure 4A: it homogeneously occupies all available space during control or below a concentration threshold, but when CA above the concentration threshold is infused the fish globally avoids this side of the setup, though some short incursions are still performed from time to time. To quantify this behavior we computed the time-based preference index (*PI*_*t*_), a quantity that has been widely used in the literature (Koide et al., 2009; Mann et al., 2003; Wakisaka et al., 2017; Lucon-Xiccato et al., 2020; Choi et al., 2021; Herrera et al., 2021) and defined as:

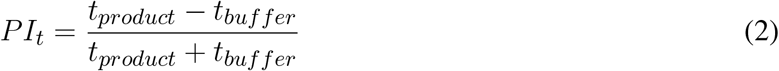

where *t*_*product*_ and *t*_*buffer*_ are the total time spent in the product and the buffer, respectively. This quantity ranges from -1 for complete repulsion to +1 for complete attraction, while *PI*_*t*_ = 0 indicates a lack of preference. The threshold concentration, defined as the concetration above which *PI*_*t*_ becomes significantly negative, was found to be close to 400μM (pH=3.5) for larvae (supplementary figure 6) and close to 100μM (pH=4.0) for juveniles (figure 4C).

**Figure 4.**
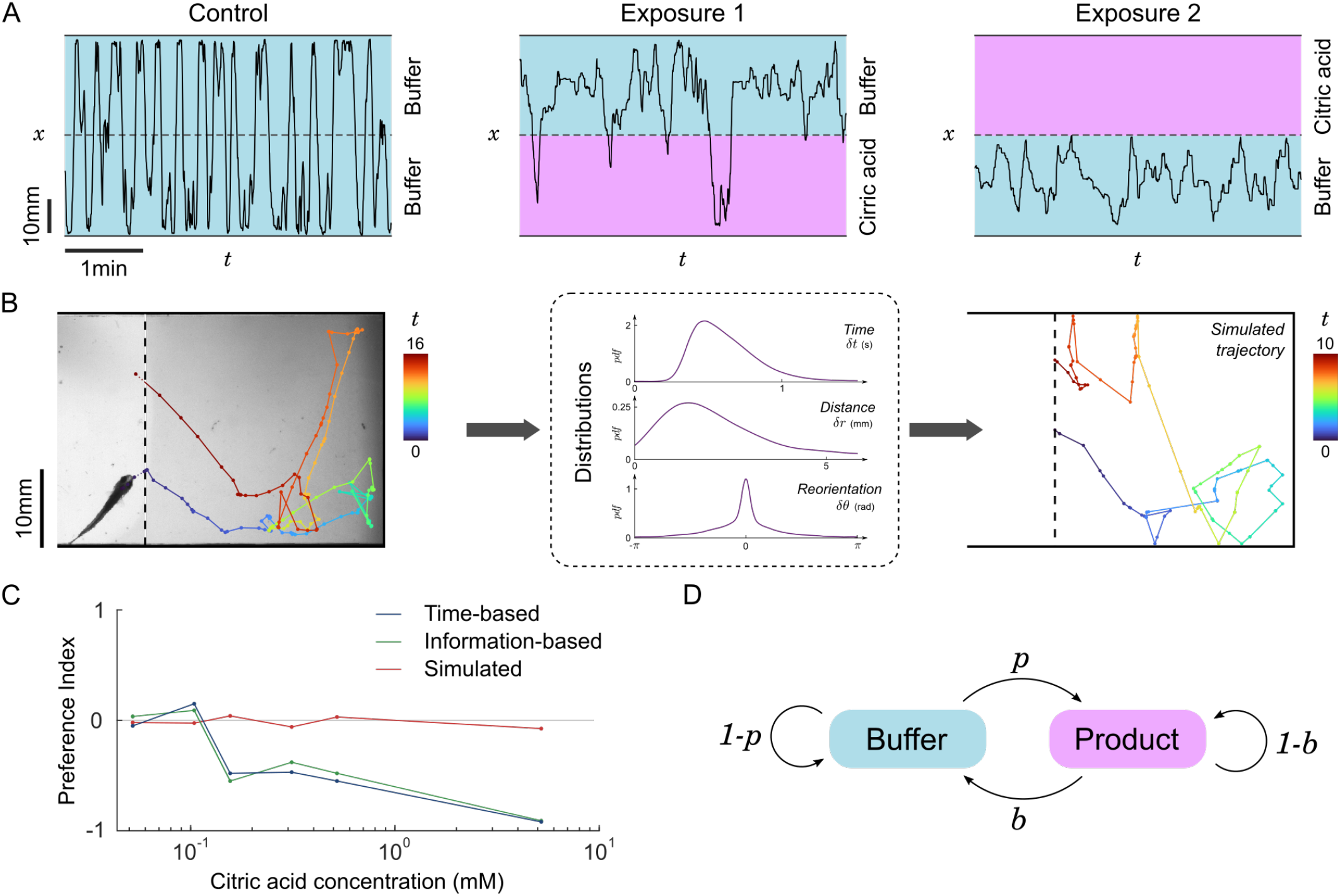
Repulsion for citric acid. **(A)** Blow-ups of lateral traces (*x*-position versus time) of the same 31 dpf zebrafish during control and two successive exposures to citric acid (476μM) on alternating sides. **(B)** Left: Sample piece of trajectory between two contacts with the interface, same fish. Color code: time (seconds). Middle: distributions of time, distance travelled and change of orientation between successive bouts from all pieces of trajectories of all fish in the same condition (product side at 476μM). Right: Example of a simulated trajectory based on these distributions. **(C)** Preference index as a function of citric acid concentration. The experimental time-based and information-based preference indices indicate a clear repulsion above a threshold close to 100μM, while the time-based preference index of simulated data remains neutral. Successive exposures are pooled together, *n*=15, 9, 6, 6, 10, 3 fish for increasing concentrations. **(D)** Two-state Markov model of transitions: animals cross the boundary from buffer to product with probability *p* and from product to buffer with probability *b*.

To test whether this repulsion is due to a modulation of some dynamical features of fish swimming in the CA region, we split the trajectories into pieces delimited by contacts with the interface as illustrated in figure 4B. We then pooled together all subtrajectories with similar conditions (buffer/product side and concentration), determined the swim bouts, and computed the distributions of times, distances, and angular reorientation between successive swim bouts (supplementary figure 7). One can use these three kinematic parameters to generate new trajectories, akin to the simulations performed in Oteiza et al. (2017) with the difference that in the present study we drew all three parameters from the distribution at every time step instead of using the average time and displacement.

The simulated subtrajectories were visually very similar to the real ones (figure 4B), but the resulting residence times (*i*.*e*. subtrajectory durations) were always statistically comparable to the control condition and the preference index calculated with the mean residence times of simulated subtrajectories remained neutral at any concentration (figure 4C).

We thus concluded that the observed excess of short trajectories in the product side was reflecting asymetric, rapid reactions of the fish upon contact with the interface. We thus focused on the outcome of the events defined by contacting the interface: upon such events, the fish had a binary choice of staying on the same side (bounce) or crossing the interface to the other side (cross). We propose a two-state Markov model of transitions (figure 4D) with two transition probabilities *p* and *b* defined as follows. The probability *p* of switching from the buffer to the product is the experimental number of events where the fish goes from the buffer to the product *n*_*BP*_ divided by all the events starting in the buffer (buffer to product and buffer to buffer) and the probability *b* of switching from the product to the buffer is the number of events where the fish goes from the product to the buffer *n*_*PB*_ divided by all the events starting in the product (product to buffer and product to product):

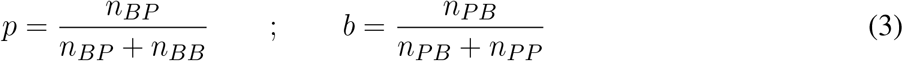

Using the above transition probabilities, the information-based preference index *PI*_*i*_ is simply defined as:

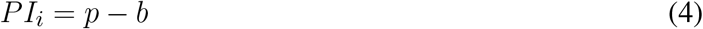

where, similarly, a value of -1 signifies pure repulsion, +1 signifies pure attraction and 0 denotes a neutral preference. The *PI*_*i*_ for citric acid as a function of concentration is shown in figure 4C, and turns out to be very similar to *PI*_*t*_.

Altogether, this suggests that the observed preference cannot be attributed to a modulation of kinematic parameters when the fish is in the product, but rather to an excess of very short subtrajectories on the product side due to frequent decisions of the fish to turn back when contacts with CA are initiated (low *p*) and to continue forward when the buffer is reached again (high *b*).

### 3.4 Changing preference to ATP

Attraction to ATP has been previously reported for adult zebrafish at concentrations above 1μM (Wakisaka et al., 2017), but we wanted to check if this was also the case for younger fish. Indeed, changes in behavioral activity in the presence of amino-acid have been reported starting from 4dpf (Lindsay and Vogt, 2004), suggesting that at least some olfactory sensory neurons and glomeruli of the olfactory bulb are already functional at this early stage.

Zebrafish larvae (4-9dpf) show no preference for ATP at 60mM but display a clear repulsion above 200mM (supplementary figure 8). This behavior being in sharp contrast with the attraction reported in Wakisaka et al. (2017) for adult zebrafish, we hypothesized the existence of a transitional period at some point during development where attraction appears.

We thus tested the preference of juvenile zebrafish (14-31dpf) for ATP between 10 and 200μM. Representative excerpts from trajectories of a 22dpf zebrafish are shown in figure 5A: the fish explored equally all the chamber during a control phase (15min), but remained essentially on the buffer side during a first exposure to ATP (15min). Throughout this period the fish had many contacts with ATP at the interface but consistently chose to stay in the buffer. During a second control period (15min) the fish resumed normal exploration of the whole chamber and, finally, during a second exposure (15min) the fish chose to stay essentially on the ATP side.

**Figure 5.**
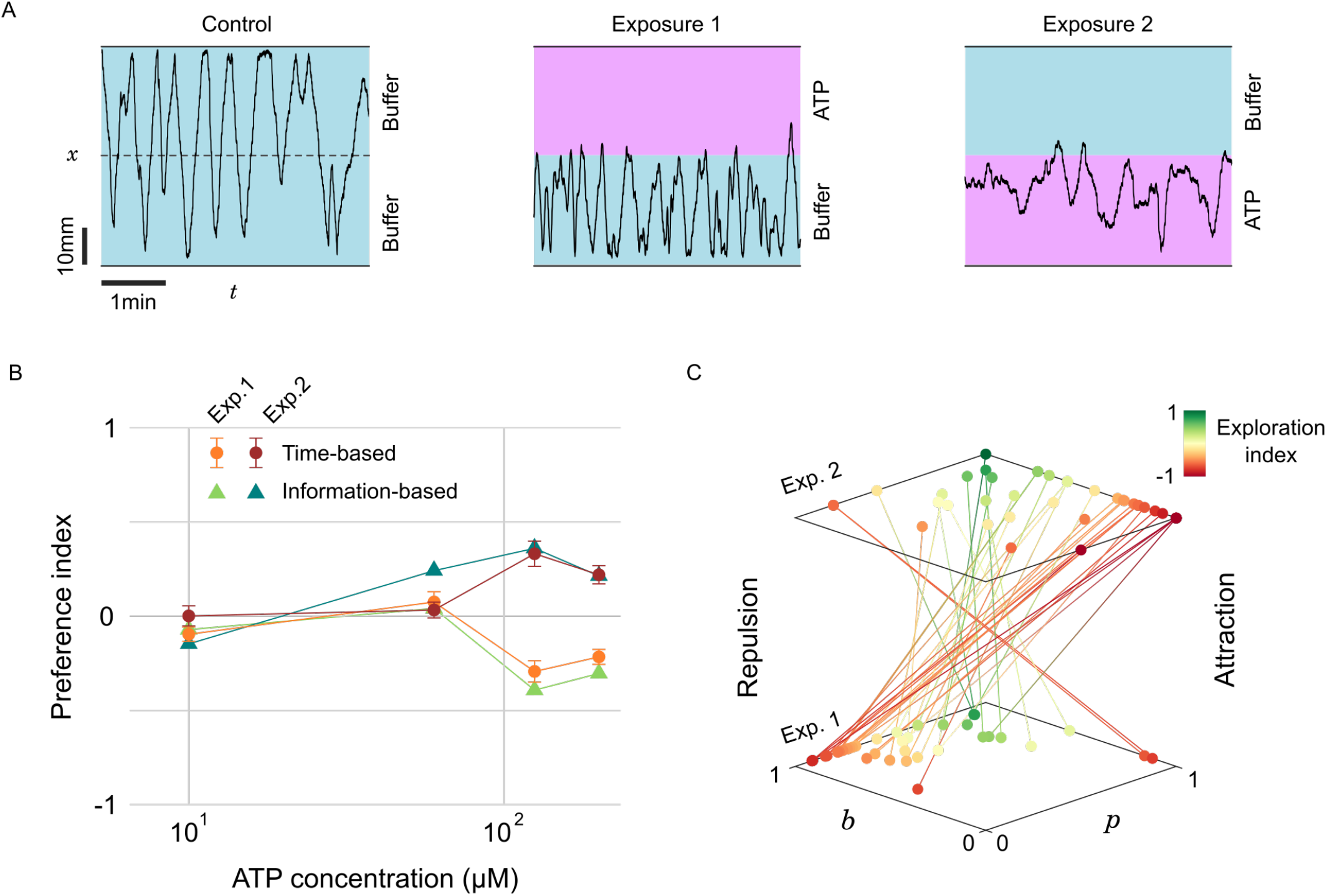
Evolving preference for ATP during successive exposures. **(A)** Blow-ups of lateral traces (*x*-position versus time) of the same 22 dpf zebrafish during control and two successive exposures to ATP (200μM) on alternating sides. **(B)** Preference index as a function of ATP concentration for two successive exposures, defined either by the residence times or by the transition statistics (*n*=28, 12, 18, 22 fish for increasing concentrations). **(C)** Transition probabilities (*p,b*) for all fish during the successive exposures to the two highest concentrations of ATP. Color encodes the exploration index, *n*=40 fish.

The *PI*_*t*_ averaged over several fish at different concentrations is displayed in figure 5C. This plot reveals a threshold at a concentration of∼100μM below which the juvenile fish did not display any preference. Above the threshold, the *PI*_*t*_ featured opposite signs for the first (repulsive) and second (attractive) exposures. As with citric acid, we quantified the probabilities *p* and *b* and computed the average information-based preference index *PI*_*i*_, which was again very similar to *PI*_*t*_ (figure 5C). The absolute *PI*_*t*_ are of moderate amplitude though, suggesting that even with a marked preference there are still, on average, frequent excursions in the less preferred zone.

We thus decided to investigate further if these excursions were homogeneously distributed among fish. Our probabilistic model allows to define for every fish another quantity of interest akin to the exploration/exploitation ratio, a familiar notion in the literature of goal-driven navigation with limited information (Vergassola et al., 2007; Barbieri et al., 2011). We define the *exploration index* as:

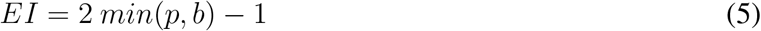

This index has a value of -1 during pure *exploitation, i*.*e*. the animal always bounces on the interface and stays on the same side, and a value of +1 for pure *exploration, i*.*e*. the animal never bounces at the interface and constantly switches side. The rationale behind this definition of the *EI* is that the fish can have a neutral preference at any exploration rate, but a marked preference can only happen in a regime dominated by exploitation.

In figure 5D we pooled all the trajectories at concentrations above the response threshold (*c >*100μM) and represented for each fish the evolution of probabilities *p* and *b* between the first and second exposures.

The color encodes the exploration index *EI*, revealing two behavioral categories: “explorative” fish (green, *n* = 8, 20%) displaying consistently high probabilities of transitions *p >* 0.5 and *b >* 0.5 for both presentations, and “exploitative” fish (red-orange, *n* = 32, 80%) that have at least one of these probabilities below 0.5, typically a low and a high probability (supplementary figure 9). For the latter category, the shift in preference is extremely salient since 29 fish (91%) displayed a clear repulsion (high *b*, low *p*) followed by a clear attraction (low *b*, high *p*). It is thus likely that the existence of some explorative fish in the dataset, upon averaging, lowers the value of the preference index.

## 4 DISCUSSION

We designed a versatile open-source millifluidic rig suitable for medium-throughput behavioral studies of exposure to chemicals. It is particularly well-adapted for testing the preference of small fish, and we used for the first time a dye to monitor simultaneously the fish and the concentration field with a single camera. The dye consisted of a diluted solution of colloids scattering light (invisible to the fish when illumination is in the infrared range), is harmless to the fish, and has a neutral hedonic value. The dye allows to quantify the extent of the mixing zone induced by the fish motion itself, but more importantly, it brings the study of trajectories to an unprecedented level of precision and paves the way to an information-based description of the behavior by focusing on the moments where the fish actually gain information over the chemical field and take decisions accordingly – which in our setup can be identified as contacts with the dye/product. Indeed, in the context of chemical sensing, such analyses have often been limited to the time-based preference index, which can be altered by long idling periods on a side due to freezing, wall-hugging (*thigmotaxis*), or reduced exploration in finite-size samples.

Our setup can be used to test virtually any organic molecule, at any concentration and multiple developmental stages. Though a relatively high throughput can be achieved (with 4 copies of the setup, a single person can manage 32 one-hour-long runs per day) as the combinatorics of possibilities is almost infinite some choices still have to be made. Innate repulsion – mostly associated with nociception – has been largely documented for algogenic and bitter compounds (Boyer et al., 2013; Candelier et al., 2015; Ohnesorge et al., 2021), for the so-called alarm substance released when a congener’s body is hurt (Speedie and Gerlai, 2008; Jesuthasan and Mathuru, 2009) and for cadaverine, putrescine and other biogenic diamines generated by decaying carcasses (Hussain et al., 2013). We quantified the repulsion to citric acid and showed that above a concentration threshold the observed general avoidance cannot be attributed to a modulation of some kinematic parameter but rather to an excess of very short subtrajectories in the product side, which is due to rapid decisions of the fish to turn back when contacts with CA are initiated.

Reports of attractive molecules are much more scarce, especially at the larval stage where sexual cues are absent and feeding capabilities are rudimental (Hara, 1992; Kasumyan and Doving, 2003). Yet, attractive chemicals are an important tool for behavioral neuroscience as it drives long-range goal-driven navigation and permits stimulus-reward association. A few promising attractive cues have been proposed for zebrafish, including kin water (Biechl et al., 2016; Gerlach et al., 2019), bile salts (Krishnan et al., 2014) as well as alanine, adenosine, and ATP (Steele et al., 1990; Wakisaka et al., 2017).

The present study unveils a developmental transition in zebrafish’s perceived valence for ATP. Wakisaka *et al*. reported ATP to be attractive for adults in the range between 1 and 50μM but to become neutral to repulsive above 100μM. Likewise, we observed repulsion for larvae appearing between 60 and 200μM, which is consistent with the hypothesis formulated by Wakisaka *et al*. that at such concentrations the activation of other sensory systems such as the trigeminal system could evoke an aversive response. However, no attraction appeared at lower concentrations suggesting that the perception of ATP was still immature. Results with juvenile zebrafish were similar to larvae at the same concentrations, with the notable difference that above the same concentration threshold the repulsion switched to attraction during repeated exposures. Using an information-based model to quantify the degree of exploration, we could separate the fish with an explorative profile – that does not display preference – from the fish actually exploiting the chemical cues. For the latter category, the change in preference between the first and second exposure was pronounced.

Several hypotheses can be formulated to interpret this inversion. First, the decrease in the nociceptive perception of highly concentrated ATP could be due to a form of habituation, though we would have expected a more gradual reversal with increasing excursions in the less preferred side, which were not observed. Experience-related functional maturation of the olfactory pathway has been reported for many vertebrates including zebrafish (Brunjes and Frazier, 1986; Kerr and Belluscio, 2006; Maher et al., 2009; Braubach et al., 2013), although generally occurring on much longer timescales. The learning of odor-reward association in adult zebrafish has been reported for timescales more comparable to our experiments (Namekawa et al., 2018).

A promising direction for future research would be to test if *Danionella Cerebrum* has a similar behavioral response to ATP at the juvenile and adult stages. As these tiny transparent fish are amenable to whole-brain functional imaging at all ages, a wide range of reward-based protocols exploiting the attraction to ATP could be envisioned to investigate the neural basis of associative learning.

## AUTHOR CONTRIBUTIONS

RC and GD designed the setup, BG and RC built the setup and performed the experiments. LP designed and produced the IR dye. BG and RC analyzed the data. All authors contributed to the manuscript.

## DATA AVAILABILITY STATEMENT

The mechanical blueprints of the setup are in supplementary figures 10-11. The electronic diagrams and data used for analyses are available upon reasonnable request to the authors. The setup software is licensed under the terms of the GNU General Public Licence. The Qt project and Arduino sketch can be found at these URLs:

- Single setup controller (integrated Raspberry 4): https://github.com/LJPZebra/dual_control/tree/arm
- Multiple (up to 4) setup control (external computer): https://github.com/LJPZebra/dual_control/tree/master

