## Supplementary information for "A scalable assay for chemical preference of small freshwater fish"

### Supplementary Material

#### Supplementary Figure 1

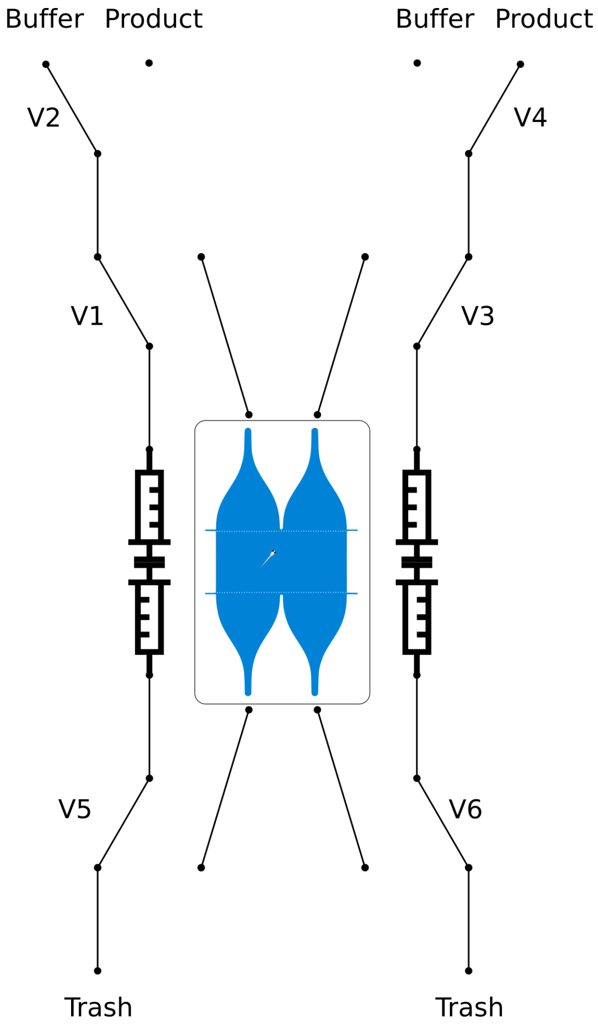


| | Injection | Aspiration Buffer | Aspiration Product Left | Aspiration Product Right |
|---|---|---|---|---|
| V1 | ■ | | | |
| V2 | | | ■ | |
| V3 | ■ | | | |
| V4 | | | | ■ |
| V5 | ■ | | | |
| V6 | ■ | | | |

**Manifold logic.** Diagram of the manifold structure with location of the six 3-way valves, and corresponding control table.

**Supplementary Figure 2**

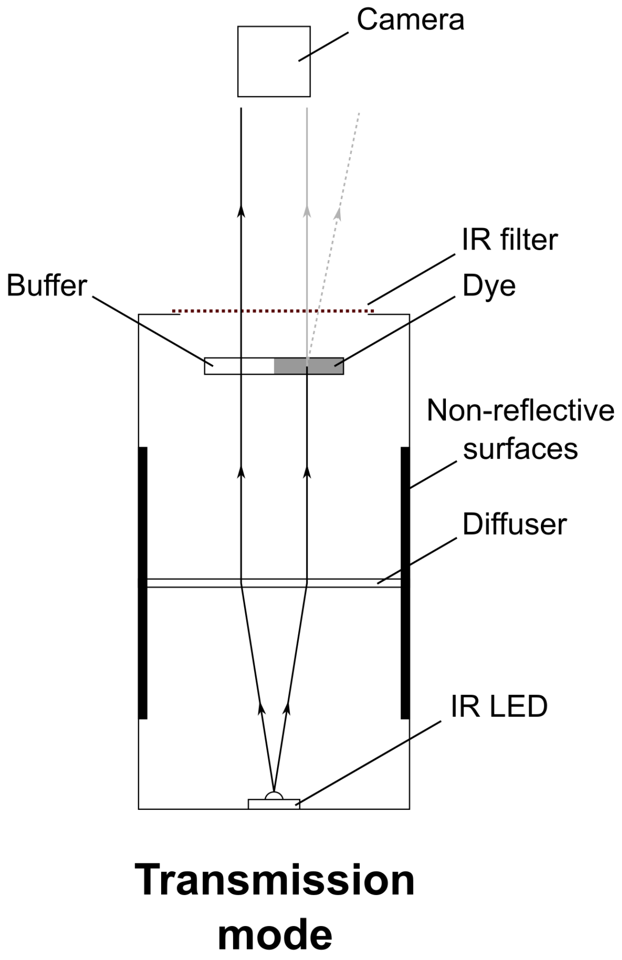


***Illumination modes***. In transmission mode (bright field) the animal and the dye appear darker than the background while in scattering mode (dark field) the animal and the dye appear lighter than the background.

**Supplementary Figure 3**

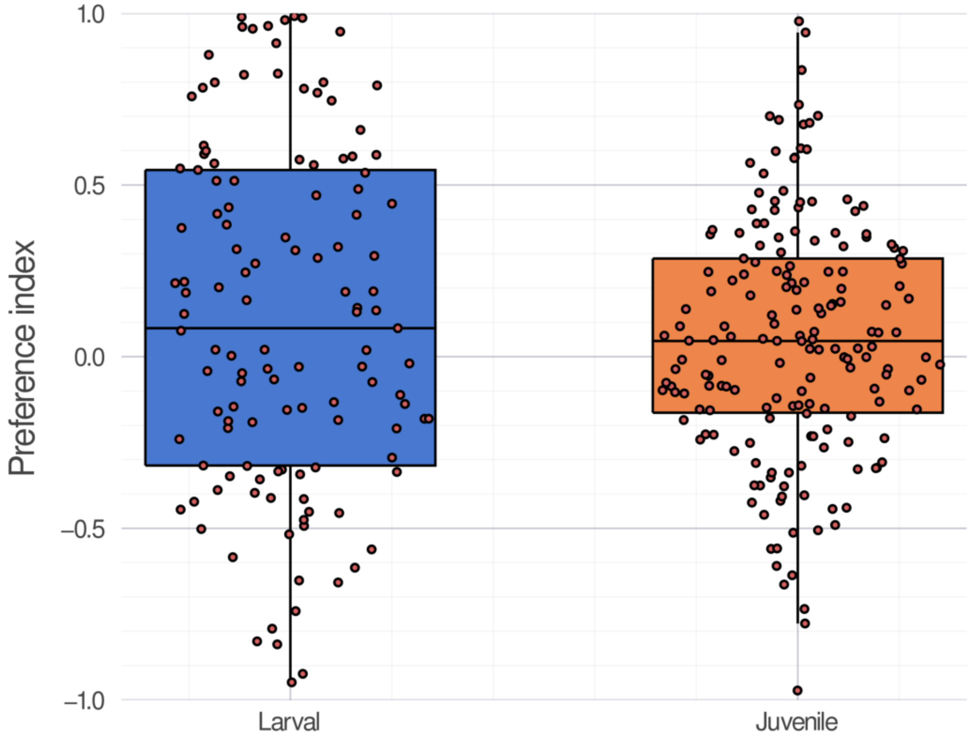


***Absence of preference for one side of the setup***. Time-based preference index of larval and juvenile zebrafish computed during the control phase of several experiments (*n*= 126 and 178, respectively). During these phases, the fish has not been in contact with the dye or any chemical.

**Supplementary Figure 4**

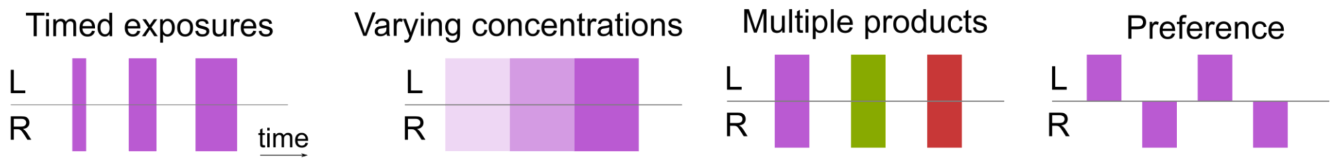


***The automated canonical protocols available in the setup***. Any combination of these protocols can be used, leading to great versatility.

**Supplementary Figure 5**

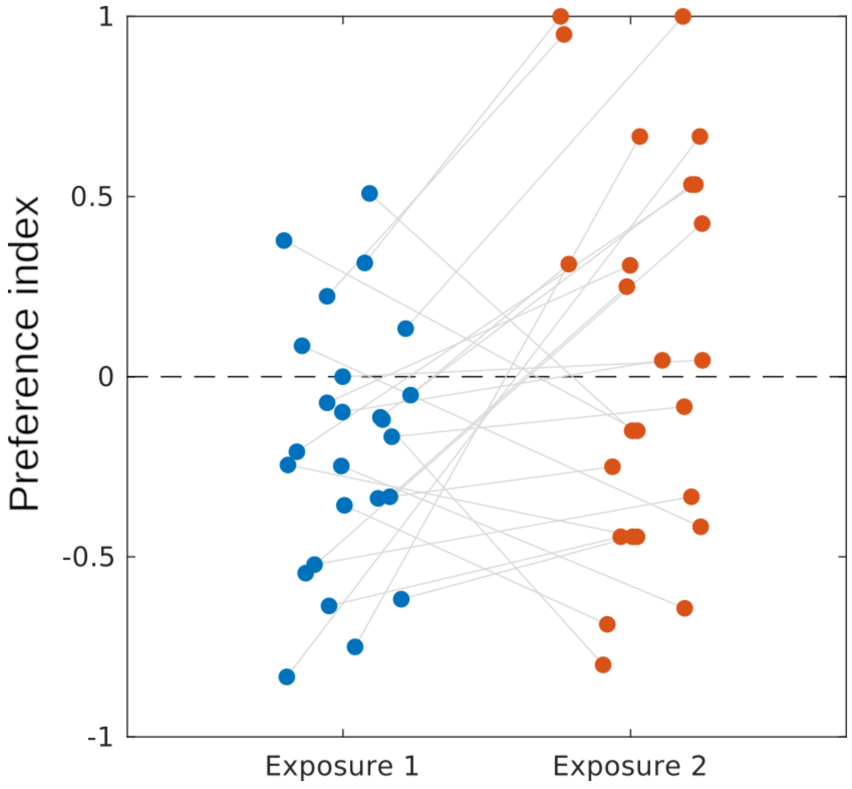


***Absence of preference for the dye***. Time-based preference index displayed for the first (blue) and second exposure (orange) of the dye (*n*=25 juvenile zebrafish). No preference bias is observed (mean preference index: -0.05) and the distributions between exposures are not statistically different (Wilcoxon signed-rank test, p>5%).

**Supplementary Figure 6**

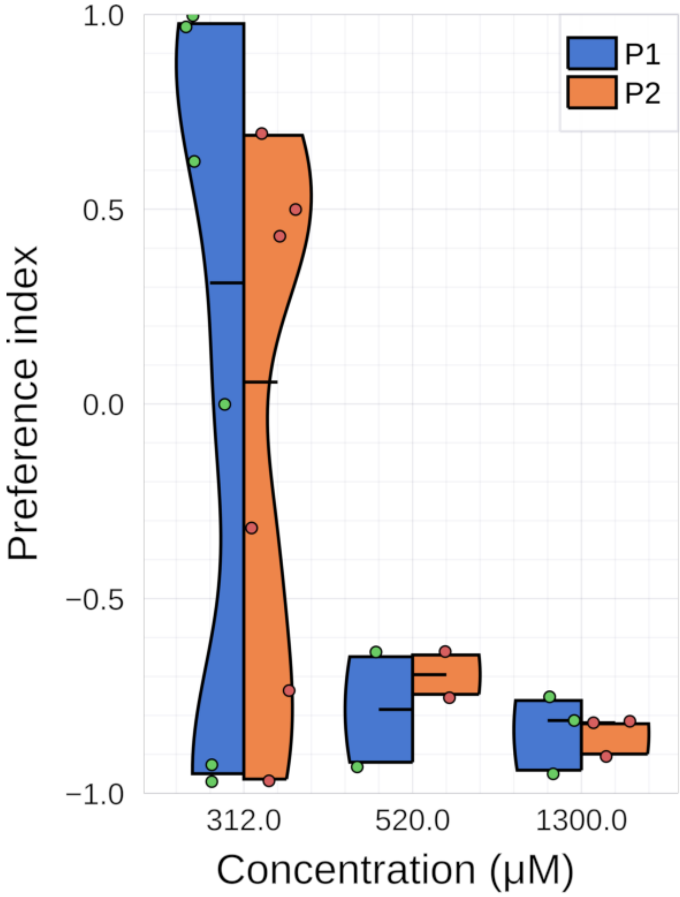


**Zebrafish larvae are repulsed by citric acid.** Time-based preference index as a function of CA concentration.

#### Supplementary Figure 7

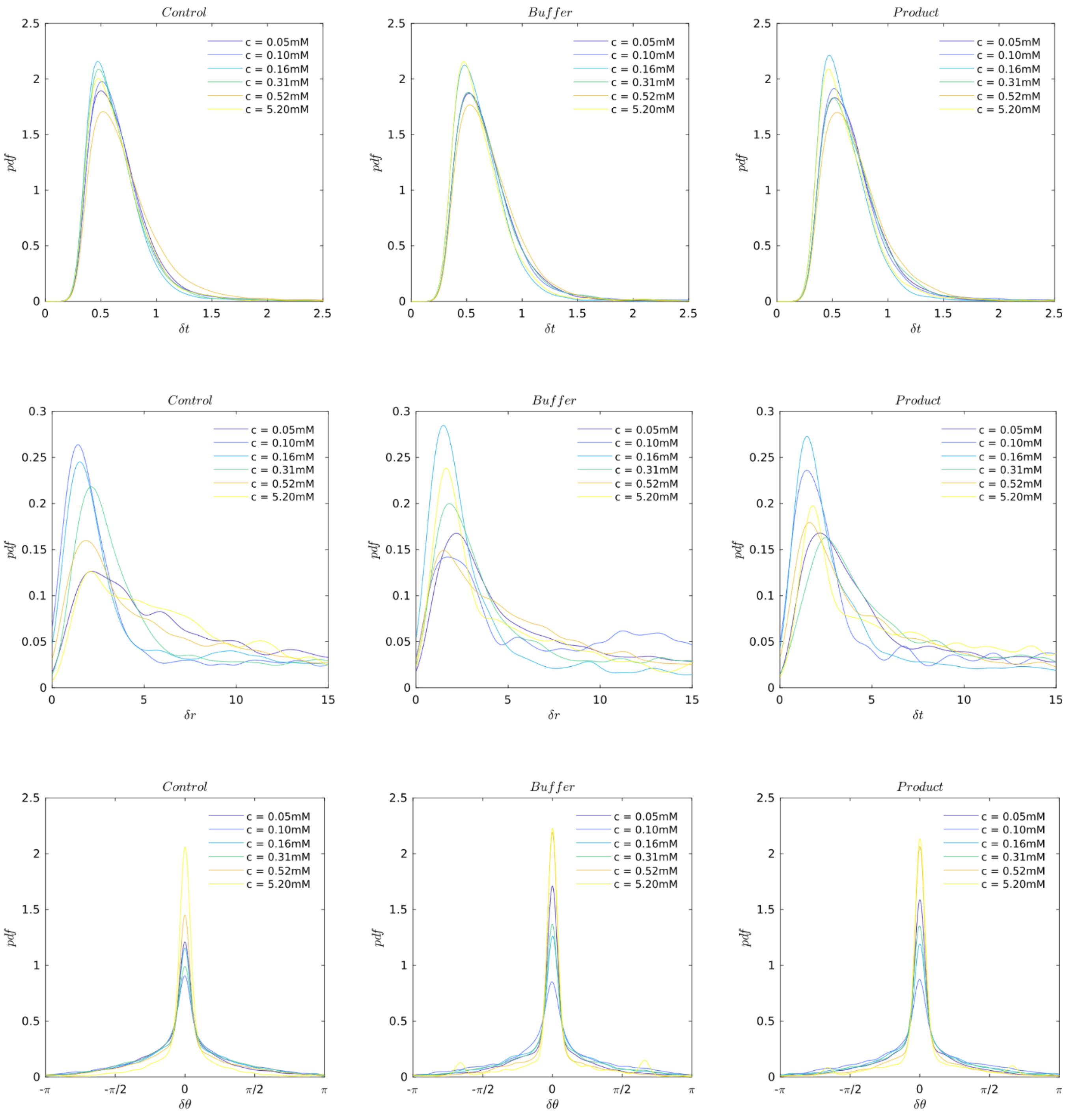


**Distributions of kinematic parameters with and without citric acid.** Distributions of times ($\delta t$, top, seconds), distances ($\delta r$, middle, millimeters) and angular reorientation ($\delta\theta$, bottom, radians) between successive swim bouts. The subtrajectories are pooled by condition: during the first 15 minutes of each trials (control) there is no dye and no product and the subtrajectories are delimited by the test chamber's middle line. During infusion, subtrajectories in the buffer and product side are isolated and pooled together. No systematic effect of the concentration can be observed, suggesting that the differences between distributions are due to the finite sample sizes.

**Supplementary Figure 8**

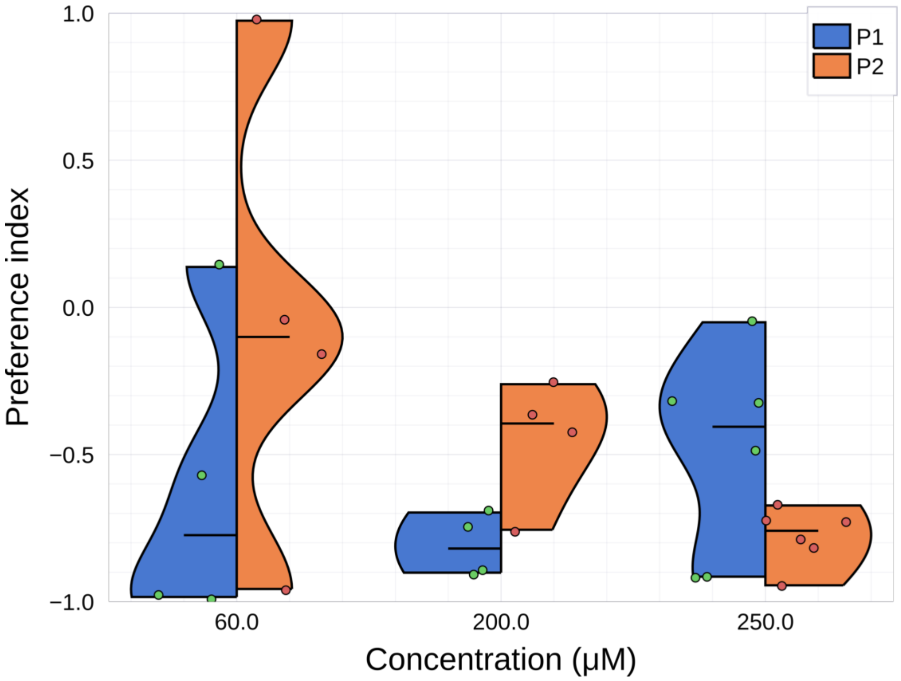


**Zebrafish larvae are repulsed by ATP.** Time-based preference index as a function of ATP concentration.

**Supplementary Figure 9**

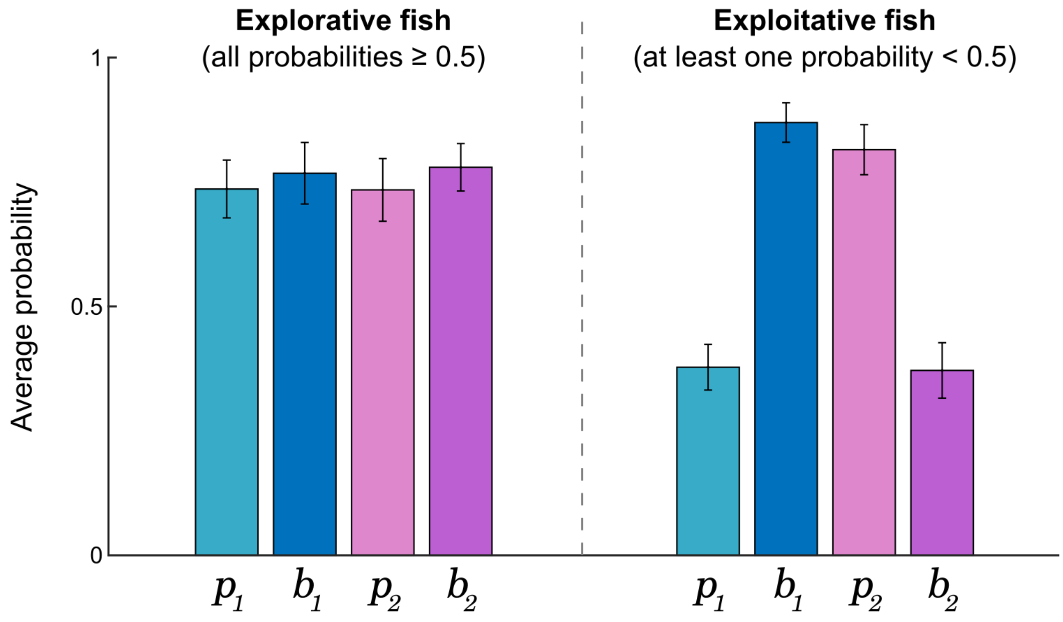


**Difference between explorative and exploitative fish.** Average probabilities $\overline{p_1}$, $\overline{b_1}$, $\overline{p_2}$ and $\overline{b_2}$ for explorative and exploitative fish. Error bars: s.e.m.

**Supplementary Figure 10**

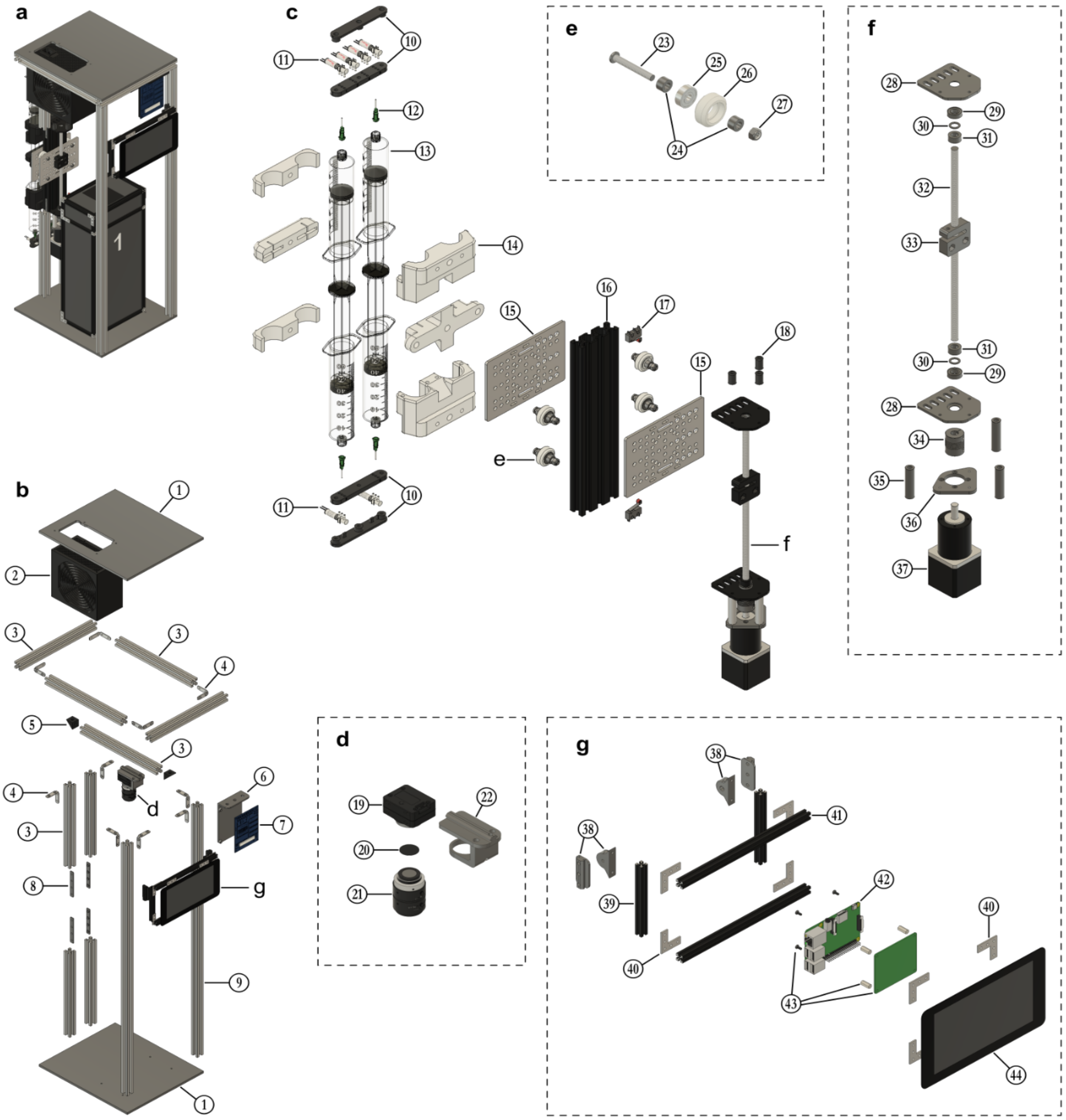

1. Laser-cut acrylic plate . x . mm (2)
2. Power supply (1)
3. Openbuilds V-Slot 20 x 20 x 250mm (9)
4. Openbuilds 90 hidden tee nut (10)
5. Openbuilds angle corner connector (2)
6. 3D printed PCB holder (1)
7. PCB / electronics (1)
8. Openbuilds quad tee nut (4)
9. Openbuilds V-Slot 20 x 20 x 750mm (2)
10. 3D printed manifolds (4)
11. 3-way microvalves (6)
12. Luer stub 18ga x 12mm (4)
13. Plastipak 60mL centered Luer lock (4)
14. 3D printed syringe holder (6)
15. Openbuilds V-Slot gantry plate (2)
16. Openbuilds V-Slot 20 x 60 x 250mm (1)
17. Roller micro switch (2)
18. 3D printed binding comb holder (3)
19. Camera - FLIR Chameleon3 (1)
20. High-pass IR filter, D=20mm (2)
21. Navitar NMV-2514 lens (1)
22. 3D printed camera holder (1)
23. Openbuilds low profile M5 screw (4)
24. 6mm spacer (8)
25. Openbuilds Xtreme solid V bearing (4)
26. Openbuilds Xtreme solid V wheel (4)
27. Openbuilds nylon insert lock nut M5 (4)
28. Openbuilds threaded rod plate (2)
29. Ball bearing 8 x 16 x 5mm (2)
30. Openbuilds shim washer 8mm (2)
31. Openbuilds lock collar 8mm (2)
32. Openbuilds 8mm lead screw (1)
33. Openbuilds anti backlash nut block (1)
34. Shaft coupler 5 x 8mm (1)
35. 40mm spacer (3)
36. Laser-cut custom adapter (1)
37. NEMA17 with 26.85:1 gearbox (1)
38. 3D printed rotating connectors (4)
39. Makerbeam 10 x 10 x 100 mm (2)
40. Makerbeam right angle bracket (8)
41. Makerbeam 10 x 10 x 200 mm (2)
42. Raspberry Pi 4 model B (1)
43. Raspberry Pi display driver kit (1)
44. Raspberry Pi LCD screen (1)

Blueprints of the general structure. **a** General view. **b-g** Exploded views of the backbone (**b**), syringe driver (**c**), camera holder (**d**), wheel assembly (**e**), translation stage (**f**) and computer and display module (**g**).

**Supplementary Figure 11**

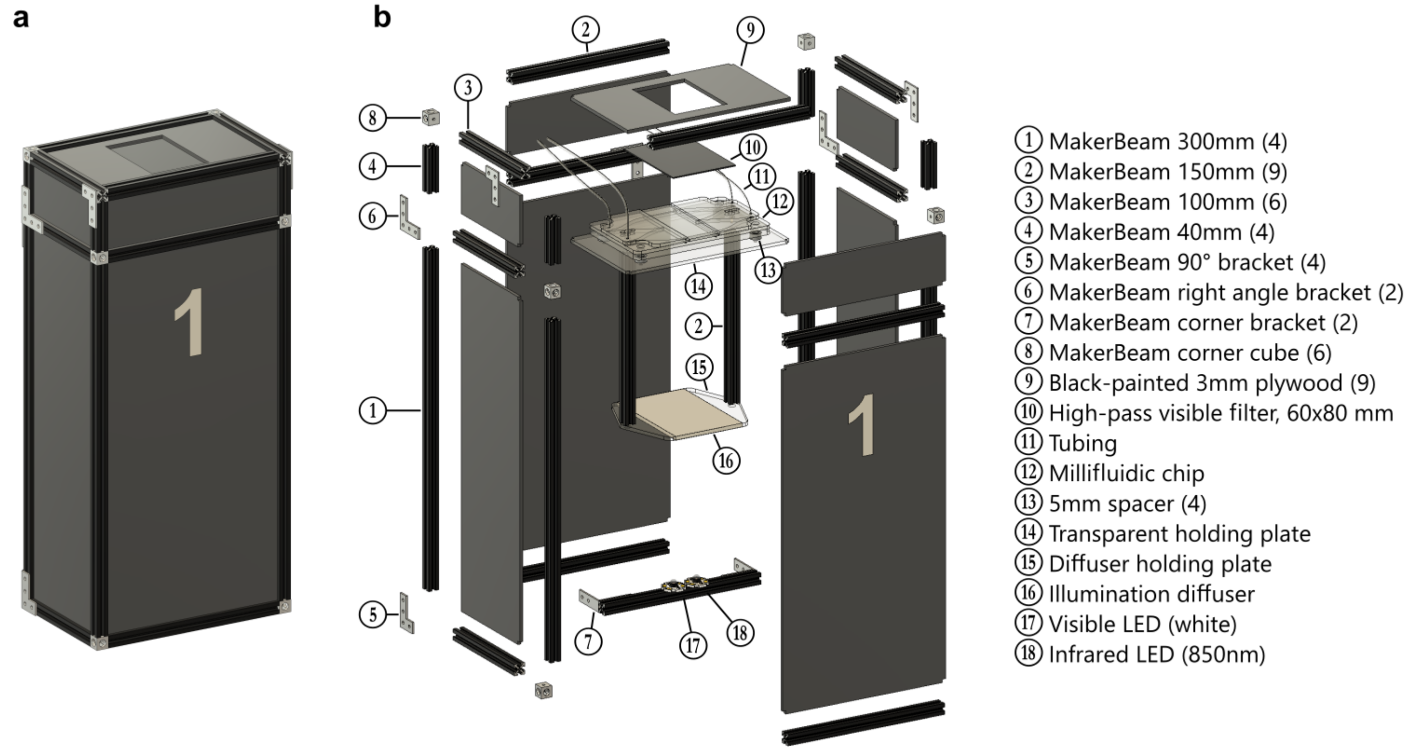


Blueprints of the illumination box and millifluidic chip. **a** General view. **b** Exploded view with parts annotation.

**Supplementary Movie 1**

***Multi-species behavioral assay.*** Compilation of short sequences of zebrafish *Danio Rerio* (larval and juvenile) *a*nd *Danionella translucida (*juvenile and adult) in the Dual behavioral assay.

**Supplementary table 1: Bill of materials**

| Category | Manufacturer | Description | Link | Qty | U.P. |
|---|---|---|---|---|---|
| Computer | Raspberry Pi | RASPBERRY PI 4 model B / 4GB | Link | 1 | 44,84 € |
| | | RPI USB-C POWER SUPPLY BLACK EU | Link | 1 | 6,52 € |
| | | RASPBERRY PI 7" TOUCH SCREEN LCD | Link | 1 | 48,92 € |
| | Corsair | Flash Voyager GTX USB 3.1 128 Go | Link | 1 | 45,57 € |
| Optics | FLIR | Chameleon3 Camera (CM3-U3-13Y3M-CS) | Link | 1 | 285,00 € |
| | | USB 3.1 Locking Cable | Link | 1 | 8,00 € |
| | Thor Labs | MVL25M23 - 25 mm EFL, f/1.4, for 2/3" C-Mount | Link | 1 | 192,80 € |
| | GoodFellow | ME303007 PMMA 0,5mm - IR transp. (80x60cm rect. + Ø20mm disk) | Link | 1 | 12,62 € |
| | Mouser | LED infrarouge 897-LZ440R608 | Link | 1 | 17,69 € |
| Electronics | Coolbox | ATX Power supply 500W Basic | Link | 1 | 20,35 € |
| | PCBWay | Custom PCB | Link | 1 | 3,93 € |
| | Arduino | Arduino nano 3.1 | Link | 1 | 16,52 € |
| | Sparkfun | Big easy driver | Link | 1 | 16,90 € |
| | | ATX Power Supply Connector - Right Angle | Link | 1 | 1,65 € |
| | | Female header | Link | 2 | 1,27 € |
| | New Japan Radio | NJM2670D2 | Link | 4 | 2,76 € |
| | Lite-on | LED through-hole green | Link | 7 | 0,09 € |
| | | LED through-hole yellow | Link | 4 | 0,09 € |
| | Bourns | Potentiometer 1 kohms | Link | 2 | 0,68 € |
| | Phoenix contact | 2-wires connector | Link | 8 | 1,29 € |
| | | 4-wires connector | Link | 1 | 2,68 € |
| | Knitter | 3-way switch | Link | 1 | 0,94 € |
| | Yageo | Resistance through-hole 560Ohms | Link | 10 | 0,09 € |
| | | Resistance through-hole 220Ohms | Link | 2 | 0,09 € |
| | | Resistance through-hole 1kOhms | Link | 1 | 0,09 € |
| Structure | Openbuilds | V-Slot® 20x20 Linear Rail (1000m) | Link | 2 | 9,04 € |
| | | V-Slot® 20x20 Linear Rail (250mm) | Link | 7 | 2,78 € |
| | | Cast Acrylic plate | Link | 2 | 7,10 € |
| | | Black Angle Corner Connector | Link | 2 | 2,53 € |
| | | Quad Tee Nut - MAKERLINK | Link | 4 | 0,76 € |
| | | 90 Hidden Tee Nut - MAKERLINK | Link | 10 | 0,85 € |
| Syringe pump | OpenBuilds | V-Slot Linear Actuator | Link | 1 | 111,88 € |
| | Phidgets | 42STH38 NEMA-17 Bipolar Stepper with 26.85:1 Gearbox | Link | 1 | 42,78 € |
| | Sparkfun | Microswitch - SPDT (Roller Lever) | Link | 2 | 1,06 € |
| | Instech | Luer stubs (LS18 - 18ga x 1/2") | Link | 6 | 0,42 € |
| | BD plastipak | 50mL syringes with Luer lock | Link | 6 | 0,44 € |
| Manifold | The Lee Company | Electrovalve LHDA0531115H | Link | 6 | 65,16 € |

| | | | | | |
|---|---|---|---|---|---|
| | VWR | Tubing (Tygon AAD04133, ID 0.05" - OD 0.09") | Link | 1 | 9,43 € |
| Box & Screen holder | Makerbeam | MakerBeam - 10x10mm aluminum profile 300mm | Link | 4 | 2,38 € |
| | | MakerBeam - 10x10mm aluminum profile 200mm | Link | 2 | 1,53 € |
| | | MakerBeam - 10x10mm aluminum profile 100mm | Link | 6 | 0,77 € |
| | | MakerBeam - 10x10mm aluminum profile 60mm | Link | 6 | 0,53 € |
| | | MakerBeam - 10x10mm aluminum profile Corner cubes | Link | 6 | 1,25 € |
| | | MakerBeam - 10x10mm aluminum profile Right angle brackets | Link | 6 | 0,58 € |
| | | MakerBeam - 10x10mm aluminum profile 90 degree brackets | Link | 6 | 0,58 € |
| | | MakerBeam - 10x10mm aluminum profile 45 degree brackets | Link | 4 | 0,58 € |
| Custom parts | - | 3D printed parts (Prusa MK3) | - | 15 | 9,00 € |
| | - | Lasercut parts (Full spectrum laser PS20) | - | 3 | 1,25 € |
| | | | | **TOTAL** | **1 430,12 €** |

**Protocol implementation**

Protocols are contained in text files that the Dual software can interpret to control the setup. Following standardized commands, one can build a custom experimental protocol by controlling the camera acquisition, valve manifolds, and infusion speed (*i.e.* the flow rate).

Here is a typical preference protocol file with alternated exposures to buffer on both sides and product on one side, followed by a quick cleaning of the setup:

```
#
# PROTOCOL: BUFFER - LEFT - BUFFER - RIGHT
#

# --- Header ------------------------------------

print:Starting protocol BRBL
data:create directory
camera:start

# --- BUFFER CYCLE ------------------------------

# --- Fill
camera:comment:Filling (buffer cycle)
switch:L:Buffer
switch:R:Buffer
switch:T:Trash
wait:5000
run:down:1500

# --- Infuse
wait:5000
camera:comment:Infusing (buffer cycle)
switch:L:Circuit
switch:R:Circuit
switch:T:Circuit
wait:5000
run:up:10000

# --- LEFT CYCLE --------------------------------

# --- Fill
wait:5000
camera:comment:Filling (left cycle)
switch:L:Stimulus
switch:R:Buffer
switch:T:Trash
wait:5000
run:down:1500

# --- Infuse
wait:5000
camera:comment:Infusing (left cycle)
switch:L:Circuit
switch:R:Circuit
switch:T:Circuit
wait:5000
```

```
run:up:10000

# --- BUFFER CYCLE ------------------------------

# --- Fill
wait:5000
camera:comment:Filling (buffer cycle)
switch:L:Buffer
switch:R:Buffer
switch:T:Trash
wait:5000
run:down:1500

# --- Infuse
wait:5000
camera:comment:Infusing (buffer cycle)
switch:L:Circuit
switch:R:Circuit
switch:T:Circuit
wait:5000
run:up:10000

# --- RIGHT CYCLE --------------------------------

# --- Fill
wait:5000
camera:comment:Filling (right cycle)
switch:L:Buffer
switch:R:Stimulus
switch:T:Trash
wait:5000
run:down:1500

# --- Infuse
wait:5000
camera:comment:Infusing (right cycle)
switch:L:Circuit
switch:R:Circuit
switch:T:Circuit
wait:5000
run:up:10000

# --- Footer ------------------------------------

wait:5000
camera:stop

# --- Clean
switch:L:Buffer
switch:R:Buffer
switch:T:Trash
wait:5000

run:down:1500:for:10000
switch:L:Circuit
switch:R:Circuit
switch:T:Circuit
```

```
wait:5000
run:up:10000

# --- Reset valves in default state
wait:5000
switch:L:Buffer
switch:R:Buffer
switch:T:Trash

print:Protocol ended
```

Here is a list of the available commands:

| Category | Command | Values | Description |
|---|---|---|---|
| Display | `print:`*`msg`* | *string* | Print the message *msg* in the user interface |
| Data | `data:create directory` | | Create a directory for storing the data of the current run. The path is automatically set with the data rooth path, the date and run number. |
| Camera | `camera:`*`state`* | *state*={start,stop} | Define the recording state |
| | `camera:comment:`*`msg`* | string | Insert a message *msg* inside the current image and append the Milestone.txt file with a line "Frame number \t TimeStamp in nanoseconds \t msg" |
| Illumination | `light:`*`type`*`:`*`status`* | *type*={VIS,IR}, *status*={ON,OFF} | Set the illumination LEDs status |
| Valves | `switch:L:`*`mode`* | *mode*={Buffer,Stimulus,Circuit} | Switch the top left syringe's mode |
| | `switch:R:`*`mode`* | *mode*={Buffer,Stimulus,Circuit} | Switch the top right syringe's mode |
| | `switch:T:`*`mode`* | *mode*={Trash,Circuit} | Switch the two bottom syringes' mode |
| Motor | `run:`*`dir`*`:`*`period`* | *dir*={up,down}, integer | Run the motor in a given direction with a velocity defined by the step period, in milliseconds |
| | `run:stop` | - | Stop the motor |
| | `run:`*`dir`*`:`*`period`*`:for:`*`time`* | *dir*={up,down}, integer, integer | Run the motor in a given direction with a velocity defined by the step period for a given duration time, in milliseconds |
| Debugging | `wait:`*`time`* | integer | Delay the execution of the next command by a given time, in millisecond |
| | `blink:`*`status`* | *status*={ON,OFF} | Set the LED status on the PCB to blink if the corresponding valve is switched |
